# Does amplitude compression help or hinder attentional neural speech tracking?

**DOI:** 10.1101/2024.02.07.578931

**Authors:** Martin Orf, Ronny Hannemann, Jonas Obleser

**Author notes:** **Correspondence should be addressed to**, Martin Orf (, Dept. of Psychology, University of Lübeck, Ratzeburger Allee 160, 23562 Lübeck, Germany), or Jonas Obleser (, Dept. of Psychology, University of Lübeck, Ratzeburger Allee 160, 23562 Lübeck, Germany).

## Abstract

Amplitude compression is an indispensable feature of contemporary audio production and especially relevant in modern hearing aids. The cortical fate of amplitude-compressed speech signals is not well-studied, however, and may yield undesired side effects: We hypothesize that compressing the amplitude envelope of continuous speech reduces neural tracking. Yet, leveraging such a ‘compression side effect’ on unwanted, distracting sounds could potentially support attentive listening if effectively reducing their neural tracking. In this study, we examined 24 young normal-hearing (NH) individuals, 19 older hearing-impaired (HI) individuals, and 12 older normal-hearing individuals. Participants were instructed to focus on one of two competing talkers while ignoring the other. Envelope compression (1:8 ratio, loudness-matched) was applied to one or both streams containing short speech repeats. Electroencephalography (EEG) allowed us to quantify the cortical response function and degree of speech tracking. With compression applied to the attended target stream, HI participants showed reduced behavioural accuracy, and compressed speech yielded generally lowered metrics of neural tracking. Importantly, we found that compressing the ignored stream resulted in a stronger neural representation of the uncompressed target speech. Our results imply that intelligent compression algorithms, with variable compression ratios applied to separated sources, could help individuals with hearing loss suppress distraction in complex multi-talker environments.

**Significant statement:** Amplitude compression, integral in contemporary audio production and hearing aids, poses an underexplored cortical challenge. Compressing the amplitude envelope of continuous speech is hypothesized to diminish neural tracking. Yet, capitalizing on this ’compression side effect’ for distracting sounds might enhance attentive listening. Studying normal-hearing (NH), older hearing-impaired (HI), and older normal hearing individuals in dual-talker scenarios, we applied envelope compression to speech streams. Both NH and HI participants showed diminished neural tracking with compression on the speech streams. Despite weaker tracking of a compressed distractor, HI individuals exhibited stronger neural representation of the concurrent target. This suggests that adaptive compression algorithms, employing variable ratios for distinct sources, could aid individuals with hearing loss in suppressing distractions in complex multi-talker environments.

## Introduction

In everyday life, people often encounter challenging hearing situations where multiple auditory signals are present. Selective attention allows listeners to prioritise a target auditory signal over distracting signals that may be occurring simultaneously (Desimone & Duncan, 1995). People with normal hearing are remarkably adept at focusing on relevant signals (even complex signals like speech) while filtering out concurrent distraction (Cherry, 1953). However, individuals with mild to moderate hearing impairments often struggle in multi-talker situations (Bronkhorst, 2000). Hearing aids are the most common treatment for hearing impairment, but even with these devices, people still face difficulties in multi-talker environments.

In recent years, computational techniques have been developed to estimate neural responses to single continuous auditory stimuli, even in the presence of other sounds (Crosse et al., 2016) Electrophysiological responses in cortical regions phase-lock to speech features in magneto/electroencephalogram recordings (Luo & Poeppel, 2007). The “temporal response function” (TRF) captures this linear relationship between continuous speech features and neural response and can be interpreted in close analogy to the classical ERP (Crosse et al., 2016; Fiedler et al., 2019). Neural phase locking to the low-frequency envelope of speech, referred to as “neural speech tracking” (Obleser & Kayser, 2019), serves as an objective measure for differentiating attended speech from concurrently ignored speech. Numerous studies have shown that individuals with normal hearing exhibit stronger neural phase locking to the envelope of attended speech compared to ignored speech (e.g., Brodbeck & Simon, 2020; Ding & Simon, 2012; Fiedler et al., 2019; Zion Golumbic et al., 2013). Additionally, there is evidence that neural phase locking to the envelope of speech correlates with speech intelligibility (Peelle et al., 2013), as well as behavioural indices of speech comprehension (Etard & Reichenbach, 2019), and that stronger speech tracking enhances trial-to-trial behavioural performance (Kraus et al., 2021).

Since neural tracking can be an objective measure for selective attention and correlates with behavioural measures, it is an interesting basis for research concerning the hearing-impaired system. However, the literature provides mixed evidence on how hearing impairment affects the neural tracking of the speech envelope. Early studies showed that poorer hearing was related to stronger tracking of the ignored envelope (Petersen et al., 2017). On the other hand, more recent studies suggest that hearing-impaired listeners show stronger neural tracking compared to the age-matched control group (Fuglsang et al., 2020). More recently, (Schmitt et al., 2022) reported enhanced speech tracking with increasing hearing loss and suggested that the hearing impaired rely more on the tracking of slow modulations in the speech signal to compensate for their hearing deficit. In contrast, other studies found no differences between older listeners with normal and impaired hearing in neural envelope tracking (Goossens et al., 2019; Presacco et al., 2019) The contradictory effects of hearing loss on neural tracking may be due to the complex interplay between ageing, the severity of hearing loss, and cognitive abilities.

Neural tracking has been shown before to depend on acoustic signal processing. For instance, vocoding can lead to delayed neural separation of competing speech during attentional selection (Kraus et al., 2021) and late cortical tracking of ignored speech is modulated differently based on signal-to-noise ratios (Fiedler et al., 2019). Furthermore, a recent study found that neural speech tracking can serve as an indicator of the benefits of hearing aid algorithms, including amplitude compression (Petersen, 2022). Overall, these findings suggest that neural speech tracking could be a useful tool for researchers seeking to understand the effects of various hearing aid algorithms, such as dynamic range compression.

Dynamic range compression (DRC) is an audio signal processing algorithm that reduces the intensity of loud sounds and effectively amplifies the contribtuon of quiet sounds. Dynamic range compression is commonly used in hearing aids to compensate for so-called loudness recruitment in hearing impaired listeners with presbyacusis and to accordingly restrain the outer-world audio dynamics into the listener’s reduced dynamic range of hearing (Kates, 2005).

However, dynamic range compression also leads to undesired side effects. For instance, compression directly affects the envelope of a speech signal. It reduces the amplitude modulation depth, alters the envelope shape, and thus alters the intelligibility-relevant temporal cues contained in speech (Stone & Moore, 1992). The envelope of speech is thus not just an acoustic feature but also affects speech comprehension (Poeppel & Assaneo, 2020; Shetty, 2016).

The rationale of the present study is as follows: We first assume that dynamic range compression impairs, in general, the neural tracking of speech, which in itself remains to be empirically shown. To this end, we measured and modelled cortical brain responses to amplitude-compressed speech in normal and hearing-impaired listeners using EEG.

Hearing aids can perform spatial signal processing, allowing them to apply different compression ratios to signals from different spatial locations (Best et al., 2021). By applying strong compression only to ignored speech, hearing aids can be potentially useful in multi-talker situations. Unlike a pure reduction of the SNR, which excessively down-regulates ignored speech (noise) and makes attention shifts more difficult, a higher compression ratio for ignored speech can strike a balance between suppression and the ability to switch speakers when needed.

We here hypothesise that amplitude compression on an ignored speech stream will increase the neural separation between the attended and ignored streams. Behaviourally, this should lead to faster response times and increased behavioural responses. The rationale here is that compression on ignored talkers will reduce their salience and corroborate the listener’s behavioural goal to attentionally suppress this stream.

To test our hypothesis, we first conducted a pilot experiment to determine the appropriate compression ratio for the following main experiments. We then recruited 24 young normal-hearing, 19 older hearing-impaired, and 12 older normal-hearing participants to participate in a target-detection paradigm in which the speech streams were either both uncompressed, both compressed, or only one of the two streams was compressed. As a control analysis, we also modelled the human auditory peripheral response (auditory nerve response and envelope following response; (Verhulst et al., 2018) to investigate the peripheral fate of compressed and uncompressed speech in normal hearing and hearing-impaired participants.

## Methods Participants

The final sample in this study consists of N=24 normal-hearing, young adults; N=19 older adults with mild to moderate, unaided hearing loss; and an age matched control group of N=12 older adults without hearing loss.

In detail, the participants in the current study were 24 young adults (18 female and 6 male), aged 18 to 34 (mean: 25.5) Each participant reported having a native language of German, having normal hearing, and having no prior neurological conditions. We measured pure tone audiometry between 250 and 4000 Hz (Figure 1) to confirm normal hearing. For the tested frequencies, all participants displayed auditory thresholds below 20 dB.

**Figure 1.**
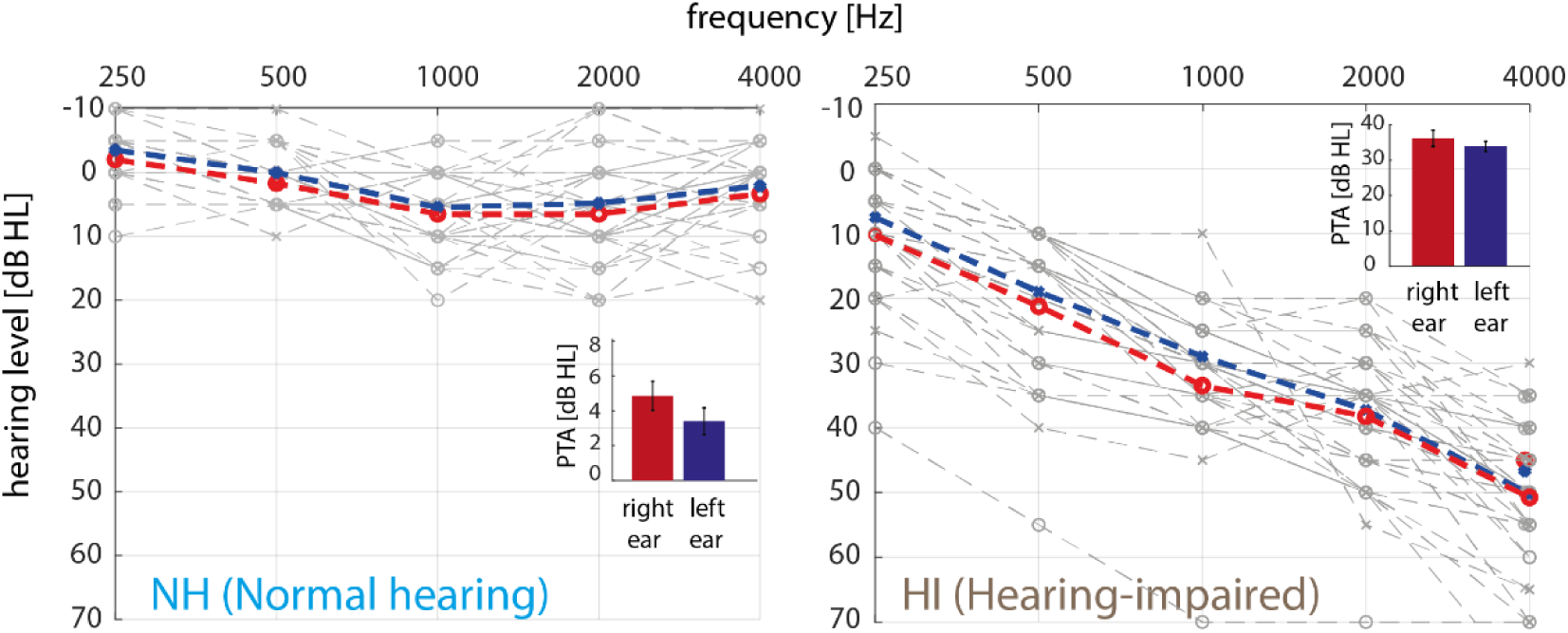
Pure tone audiometry. Left: Normal hearing participants (NH, N = 24). Right: Hearing-impaired participants (HI). Average pure tone hearing thresholds of the left (blue cross) and right (red circle) ear across frequencies (250 -4000 Hz) for N = 19 hearing-impaired participants. Grey lines show single-subject hearing thresholds (left-cross; right-circle). Inset shows pure tone average (PTA); bars indicate the mean ± standard error, grey dots indicate single-subject data.

We also enrolled hearing impaired participants between the ages of 50 and 75 with mild to moderate presbycusis, defined as a pure tone average (PTA) between 20-50 dB HL (Humes et al., 2012) similar hearing thresholds in both ears with a maximum difference of 10 dB, and no or less previous experience with compression in hearing aids, either unaided or with no longer than one year of prior use. Of the originally recruited 21 participants with hearing loss, three of them had to be excluded from the analysis (for data loss during EEG recording; very poor task performance; and hearing loss of non-presbycusis aetiology, respectively). The remaining N = 19 participants were aged between 57 to 75 years old, with an average age of 66.8 years.). To assess their hearing ability, pure tone audiometry was also performed for frequencies ranging from 250 to 4000 Hz (Figure 1). All hearing-impaired participants showed the typical sloping progression to higher frequencies in auditory thresholds, with pure tone averages ranging from 25-42 dB HL and an average of 36 right and 34 left dB HL.

To have an age-matched control group, we secured an additional of N=12 older participants aged between 52 to 73 years old (average age of 65 years) without hearing impairment (PTA < 25 dB HL, average PTA∼ 15 dB HL, Fig.7 A).

All participants provided written, fully informed consent and were paid 10 euros per hour. The study was approved by the local ethics committee of the University of Lübeck.

### Stimulus materials

We presented audio versions of two different narrated book texts, “Ludwig van Beethoven Basiswissen” and “Sophie und Hans Scholl Basiswissen”, both of which had been pre-recorded by professional talkers (male and female speaker). We selected audio streams that had not previously undergone amplitude compression. The two audio streams overlapped in time but were spatially separated (see “Experimental Setup” below) and were presented at an average intensity (SPL mixture) of about 65 dB(A), which is comparable to the volume of a normal conversation.

Using customised MATLAB code, the stimuli were processed in the following steps (Version 2018a Mathworks Inc., Natick, MA, United States). The audio files had a 44.1 kHz sampling rate and a 16-bit resolution. The maximum duration of silent periods was reduced to 500 ms (O’Sullivan et al., 2015).

By selecting 400 ms of the original audio stream and repeating it immediately after, we added brief repeats to both audio streams (Marinato & Baldauf, 2019; Orf et al., 2023). At least two seconds after the stimulus began, the first repeat was shown. By linear ramping and cross-fading, each repeat was incorporated into the sound stream. Utilizing a window of 220 samples (5 ms) from the down ramp’s end and the first 220 samples (5 ms) from the repeat itself, linear ramping was performed (up ramp). The cross-fading was accomplished by combining the up and down ramps.

In order to prevent undetectable repeats of weak sound intensity, we further used an rms (root mean square) criterion, which required that the repeat’s rms be at least equal to the rms of the stream from which it was drawn (Orf et al., 2023).

Using a digital dynamic range compressor built into MATLAB, we applied amplitude compression to the speech streams (Giannoulis et al., 2012). We set the compressor parameter as follows: Attack time = 2 ms; release time = 15 ms; threshold = –40 dB. Note that we for this proof-of-principle study we used a higher compression ratio and faster attack and release times compared to standard hearing aid processing (e.g., Kates, 2005). To avoid clipping, both the uncompressed and compressed audiobooks were peak limited to 95% full scale. The compression ratio was determined by a pilot experiment (Figure 2A).

**Figure 2.**
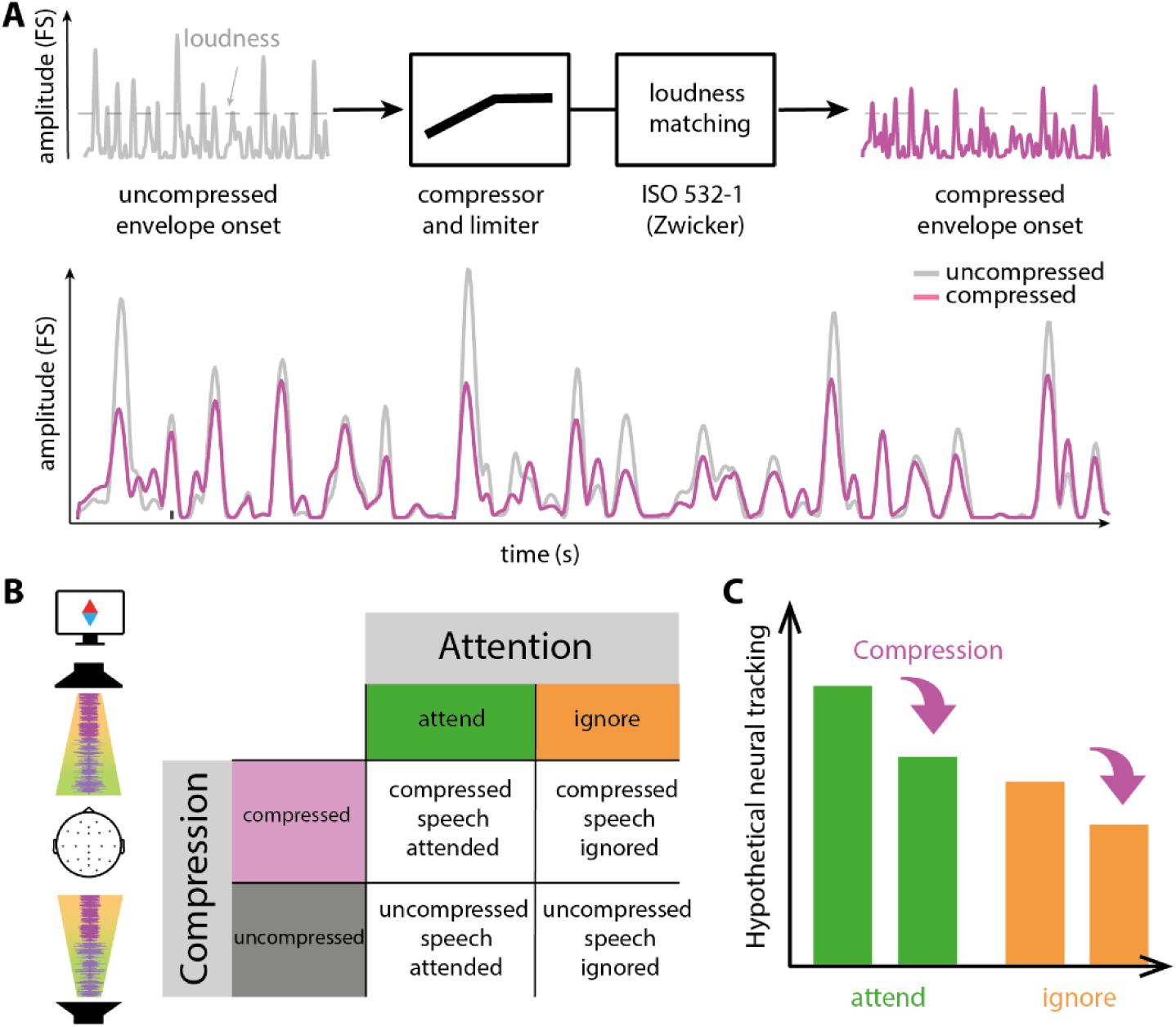
Stimulus processing, experimental design and hypothetical results. The most important stimulus processing flow is shown by **A**. The stimulus is expressed as the envelope onset of the speech signal. First, the signal is processed by the compressor (ratio: 1/8, attack time: 2 ms; release time: 15 ms; threshold: - 40 dB). Importantly, the limiter was then applied to both signals the uncompressed and compressed signals- to avoid clipping. The compressed audio segments were matched to the loudness of their uncompressed counterparts by using a MATLAB algorithm based on Zwicker **B**. Left: Experimental setup. Two loudspeakers are placed in front (0°) and back (180°) of the participant. Speech streams are simultaneously presented over both loudspeakers. A screen was placed in front of the participant. A spatial cue indicated to which location participant had to attend. Right: Experimental paradigm. We had a quasi-factorial design with the factors attention (2- levels: attended and ignored) and compression (2-levels: compressed and uncompressed). Importantly, attending and ignoring always happened at the same time, while the factor compression was fully balanced. **C** Hypothetical results. For the main effects, we would expect that attention (to-be-attend to the cued stream) has a positive effect on the dependent variables, which means that attention leads to increased behavioural and neural results. In contrast, compression would have a negative effect on the dependent variables, that is, decreased neural and behavioural results.

For pairing a-to-be attended and a to-be-ignored audio segment, we created four pairs of audio segments for the: uncompressed & uncompressed, uncompressed & compressed, compressed & uncompressed, and compressed & compressed; to maintain a balance between compressed and uncompressed segments in the two streams. Each pairing had a duration of 5 minutes. The pairings were arranged in a balanced, pseudo-random manner (Figure 2B). Due to the structure of our experiment, attending and ignoring occur simultaneously. To ensure consistency and accuracy in our analysis, we investigated only those trials where attention and compression were applied concurrently. Moving forward, we will refer to these trials as “attentional pairs.”

The root-mean-square (RMS) is frequently used to match the intensity of distinct audiobooks. However, it was demonstrated that the perceived loudness of RMS matched uncompressed and compressed speech differs (B. C. J. Moore et al., 2003) Here, we matched the perceived loudness for time-varying acoustic signals based on Zwicker using an internal MATLAB function (Zwicker & Scharf, 1965)

We conducted a psychophysical experiment to confirm the viability of this algorithm using the stimuli we had previously used. The results suggest no significant differences in loudness perception between uncompressed and Zwicker matched compressed speech.

The cue was presented at the centre of the screen (1920×1080 resolution, Wimaxit 15.6-inch Portable HDMI Screen) in front of the participant (distance: 1 m). The spatial cue consisted of two triangles which had a size of 1.3° visual angle pointing to the front and back sound sources the two triangles had different colours blue and red. Participants had to attend either to the red or the blue triangle. Since the cue and the fixation cross were presented at the same time as the auditory stimuli, we ensured that the possible interference between visual and auditory neural responses was as small as possible. In order to achieve this, the cue was linearly faded in and out (50 ms each) to create a seamless transition between the fixation cross and cue.

### Experimental Setup

The experiment was conducted in a soundproof chamber with two loudspeakers (Speaker 8020D, Genelec, Denmark) placed at a one-meter radius in the front and back, respectively. The speakers were placed 1.20 meters above floor, and a chair was positioned in the centre of the radial speaker array, facing the loudspeaker at position 0° in the azimuth plane (Figure 2B). Participants received a briefing on the experiment in advance. Each participant was instructed to keep their eyes open, keep their gaze on the center of the screen, and sit as comfortably as they could. A chin rest was utilized to prevent head movement. Each participant had their chin rest adjusted in height.

### Experimental Procedure

To study amplitude compression in a competing-talker paradigm, we developed a new experimental procedure. The experiment was created using Psychophysics Toolbox extensions (Brainard, 1997) and MATLAB (MathWorks, Natick, MA, USA). Two audio streams were played simultaneously for participants. Each trial started with a cue that specified which stream to attend, displayed for 500 milliseconds. A fixation cross was then shown for the remainder of the trial (19.5 s), while the auditory stimuli continued to play in the background. Trials were presented continuously, with the next trial starting immediately after the previous one ended.

Participants were required to identify short repeats (*Original sentence:* “The house is nice.”; *With short repeat:* “The house house is nice.”) in the target stream, with six repeats included in each trial and randomly divided between the two streams. Prior to data collection, the experiment was explained to participants, emphasising the importance of listening to the target stream’s content and responding as quickly and accurately as possible to a repeat. Participants were given a single sentence containing one repeat to acquaint them with the repeats and were asked to provide oral feedback if they were able to recognise it. Additionally, participants completed six practice trials that were identical to the main experiment but used different audio streams. The main experiment lasted approximately one hour and included 196 consecutive trials split into four blocks, with participants having the opportunity to rest between each block. All participants performed the experiment without using hearing aids.

### Sound pressure level adjustments for hearing-impaired participants

The hearing-impaired participants completed the same experiment as their normal hearing counterparts, using the same stimuli but with different randomizations of conditions. However, we made one change: We adjusted the overall sound pressure level of the experiment based on each participant’s hearing loss. To determine their hearing threshold, we used 500 ms parts of the stimuli that were presented in the experiment itself over free-field loudspeakers. We employed a combination of limit and constant stimuli methods to establish the threshold for the experimental stimuli. First, we presented pairs of compressed and uncompressed stimuli snippets (one over the front loudspeaker, one over the back), with one always louder than the other. In 3 dB steps, we decreased the sound pressure level of the signals each time the participant pressed a button to indicate they could hear the sound snippet. Once the participant stopped responding, we set this level as a reference for the method of constant stimuli. We then presented three different levels in 2 dB steps before and after the reference level, with each level presented 10 times in random order for a total of 70 presentations. We used the participants’ responses to fit a psychometric function and obtained the SRT50 of this function as the new determined threshold. We added 35 dB to this threshold to set the presentation level. However, we asked each participant after the procedure if the overall presentation level was appropriate for them. If they did not agree, we adjusted the presentation level in 5 dB steps until it matched their reported most comfortable perceived loudness. On average, the presentation level was 72 dB ranging from 65 to 88 dB SPL.

### Data acquisition and pre-processing

A 24 electrode EEG-cap (Easycap, Herrsching, Germany; Ag-AgCl electrodes positioned in accordance with the 10-20 International System) connected to a SMARTING amp was used to record the EEG (mBrainTrain, Belgrade, Serbia). The portable EEG system sends the signal via Bluetooth to a computer for recording. Using the program Smarting Streamer (mBrainTrain, version: 3.4.2), EEG activity was captured at a sample rate of 500 Hz. Impedances were kept under 20 k**Ω** while impedances were used as an online reference during recording using electrode FCz. The Fieldtrip-toolbox, built-in functions, and MATLAB (Version 2018a Mathworks Inc., Natick, MA, United States) were used for offline EEG preprocessing (Oostenveld et al., 2011). High-and low-pass filters were applied to the EEG data between 1 and 100 Hz, and the electrodes M1 and M2 (the left and right mastoids) were averaged (two-pass Hamming window, FIR, filter order: 3f_s_/f_c_). An anti-aliasing filter was applied prior to these steps to prevent aliasing artifacts. On the EEG data from every participant, an independent component analysis (ICA) was performed.

Prior to ICA, M1 and M2 were removed. Visual inspection was used to identify and remove ICA components associated with eye blinks, eye movement, muscle noise, channel noise, and line noise. On average, 7.89 out of 22 components (SD = 2.74), were disqualified. Back projected to the data were elements not connected to artifacts. Clean EEG data were processed further. Frequencies up to 8 Hz are associated with neural speech tracking (Luo & Poeppel, 2007). EEG data were therefore low-pass filtered once more at 10 Hz (two-pass Hamming window, FIR). EEG data were then segmented into epochs that matched the trial length of 20s and resampled to 125 Hz.

### Extracting the speech envelope

By calculating the onset envelope of each audio stream, the temporal fluctuations of speech were quantified (Fiedler et al., 2017). First, we used the NSL toolbox (Chi et al., 2005) to compute an auditory spectrogram (128 sub-band envelopes logarithmically spaced between 90 and 4000 Hz).Second, to create a broad-band representation of the temporal envelope, the auditory spectrogram was averaged across frequencies. Third, the half-wave rectified first derivative of the onset envelope was obtained by computing the first derivative of this envelope and zeroing negative values. In order to match the EEG analysis’s target sampling rate, the onset envelope was lastly down sampled (125 Hz). By using the onset envelope instead of the envelope, the envelope is moved in time. It’s notable that the TRF obtained by using the onset envelope as a regressor resembles a conventional ERP more than when the envelope is used as the regressor (Fiedler et al., 2017).

### Temporal response function and neural tracking estimation

A temporal response function (TRF) is a condensed brain model that illustrates how the brain would process a stimulus feature to produce the recorded EEG signal if it were a linear filter. To calculate the TRF, we employed a multiple linear regression method (Crosse et al., 2016). In order to more precisely predict the recorded EEG response, we trained a forward model using the onset envelopes of the attended and ignored streams (e.g., Fiedler et al., 2019) In this framework, we examined delays between envelope changes and brain responses of between 0 and 500 ms.

To address EEG variance related to processing behaviourally relevant repeats and corresponding evoked responses, we added all repeat onsets and button presses as nuisance regressors using stick functions. A stick function is a vector used in modelling, where non-event times are represented by zeros and events, such as the onset of repeats or button presses, are marked with a value of 1. These repeat onsets were added independently of the speech envelope regressors and chosen almost randomly (within SNR threshold) for each speech stream.

To prevent overfitting, we used ridge regression to estimate the TRF and determined the optimal ridge parameter through leave-one-out cross-validation for each participant. We predefined a range of ridge values, calculated a separate model for each value, and averaged over trials to predict the neural response for each test set. The ridge parameter with the lowest mean squared error (MSE) was selected as the optimal value specific to each subject. TRFs were estimated from trials in the experiment. To avoid cue conflicts, the first second of each trial was excluded. One model was trained on 192 trials using predictor variables for the onset envelopes of attended and ignored streams, as well as stick functions for repeats and button presses. These were modelled jointly (same regressor matrix) using the same regularization.

Neural tracking measures the representation of a single stream in the EEG signal, using TRFs to predict the EEG response. By using Pearson correlation to compare the predicted and actual EEG responses, the neural tracking (r) was calculated. By using the leave-one-out cross-validation method, we were able to predict the EEG signal on single trials (see above). A sliding-time window (48ms, 6 samples, 24ms overlap) calculated neural tracking accuracy over TRF time lags, resulting in a time-resolved neural tracking (Fiedler et al., 2019; Hausfeld et al., 2018; Kraus et al., 2021; O’Sullivan et al., 2015).

### Peripheral auditory modelling

We employed a computational model of the human auditory periphery developed by Verhulst and colleagues (2018) to simulate model outputs to the speech signals used in our experiments presented at 65 dB SPL. We randomly selected 100 speech snippets from our uncompressed stimulus material and applied the same processing pipeline (including compression and loudness matching) as used in our main experiment to create a set of 100 compressed speech snippets and a corresponding set of 100 uncompressed snippets. We modelled the firing rate of the auditory nerve (AN) and the envelope following response (EFR) for both normal hearing and hearing-impaired participants, simulating a typical mild to-moderate presbycusis (hearing loss due to aging) starting at 1 kHz and sloping to 35 dB HL at 8 kHz. As the AN response varies with frequency, we focused on four center frequencies (500, 1000, 2000, and 4000 Hz) that are particularly relevant to speech in audiology (Sweetow & Silverman, 1994). The EFR, which reflects the neural processing of the temporal envelope, was modelled without frequency dependence.

To analyse the output of the auditory nerve (AN) in greater detail, we employed a mixed model. The dependent variable was the log-transformed spike rate of the modelled AN. We used the same speech snippet to generate both uncompressed and compressed outputs, for both normal hearing (NH) and hearing-impaired (HI) conditions, resulting in four different AN output for each speech snippet. To account for the quasi-repeated measures nature of the data, we included speech snippet as a random effect in the mixed model. In addition to hearing impairment (NH, HI) and signal manipulation (uncompressed, compressed), we also included frequency (500, 1000, 2000, and 4000 Hz) as a factor in the model.

### Statistical analysis

We employed various statistical methods to address our research questions. In our pilot study, which involved 6 participants, we employed a double bootstrapping technique to obtain more accurate confidence intervals for our estimates. This method involved two levels of bootstrapping: First, we generated a large number of 2,000 bootstrap samples from our data and computed the statistic of interest for each sample. In the second step, a smaller number of 200 additional bootstrap samples were drawn from the distribution of the first level’s bootstrap estimates. This iterative process allowed us to calibrate the confidence intervals, reducing bias and improving the accuracy of our estimates in this initial phase of the research (Penn, 2020).

To examine the behavioural data in relation to detected repeats, we utilised logistic regression to model the binary outcome (hit = 1/miss = 0) of each repeat. We used a mixed model to predict the continuous dependent variable response speed (1/response time). We incorporated both attention (attend/ignore) and compression(compressed/uncompressed) categorical predictors in both models to examine their main effects and interactions measured. To assess statistical differences in neural tracking, we employed mixed models with the same categorical predictors as previously mentioned. However, the difference was that we utilised the models to predict neural tracking. Additionally, we included the categorical predictor space (front/back) to control the spatial assignment of loudspeakers in the setup for all models. In pseudo-code, the formula for the linear model can be expressed as *neural_tracking ∼ 1 + Attention + Space + Compression + PTA + age + Attention:Space + Attention:Compression + Space:Compression + Attention:Space:Compression+( 1 | Subject_id )+( 1 | Trial )*.

For the comparisons within attentional pairs, we used paired t-tests. We used jamovi for, gamlj package in R for fitting generalized linear mixed models (Jamovi Project, 2020), and MATLAB’s fitlme function for fitting linear mixed models (Math-Works Inc., 2020).

### Statistical analysis on time series

We investigated whether there were differences in time points in time-resolved neural tracking and TRF between subjects in different conditions and attentional pairs. To do this, we utilized a two-level statistical analysis known as cluster permutation test, which was implemented in Fieldtrip (Oostenveld et al., 2011). The analysis was conducted on data from 22 channels. At the single-subject level, we performed one sample t-tests to assess TRF and time-resolved neural tracking differences. Clusters were defined based on resulting t-values and a threshold set at p < 0.05 for at least three neighbouring electrodes. The observed clusters were compared to 5,000 randomly generated clusters through a permutation distribution using the Monte Carlo method to correct for multiple comparisons (Maris & Oostenveld, 2007). The cluster p-value was determined by the relative number of Monte Carlo iterations in which the summed t-statistic of the observed cluster exceeded that of the random clusters.

## Results

### Pilot study: the effects of compression ratio on neural tracking of speech

In this pilot experiment, six young individuals with normal hearing were asked to listen to a narrative story that contained randomly balanced parts of three different compression and expansion ratios. The compression ratios were 1:2 and 1:8, while the expansion ratio was 2:1. The participants were also presented with a baseline condition where the narrative story was uncompressed. The task was to listen to the content of the presented narrative story. The objective of this pilot experiment was to identify an appropriate compression ratio to be used in a follow-up study.

For this pilot study, we opted to use a decoding approach rather than an encoding approach. The main reason for this choice was to take advantage of the higher accuracy that can be achieved by using all EEG channels for reconstruction also given the small sample. Additionally, we were able to avoid potential confounds related to the temporal response functions of the brain to repeated stimuli, as well as confounds related to button presses since repeats were not included in this pilot experiment.

Figure 3A depicts the averaged decoding accuracy for different compression and expansion ratios. To investigate differences between conditions we used bootstrapped CIs. For the comparison of compression ratio 1:2 and expansion ratio 2:1 versus uncompressed baseline (raw) the bootstrapped CI included zero, indicating a non-significant difference. However, for the difference between the 1:8 compression ratio and uncompressed speech, the bootstrapped confidence interval (CI) did not include zero, indicating a significant difference. Specifically, the decoding accuracy for the 1:8 compression ratio was significantly lower than that for the uncompressed speech, as the entire CI was below zero. Interestingly, each participant shows a decreased decoding accuracy for the 1:8 compressed speech signal (Figure 3B).

**Figure 3.**
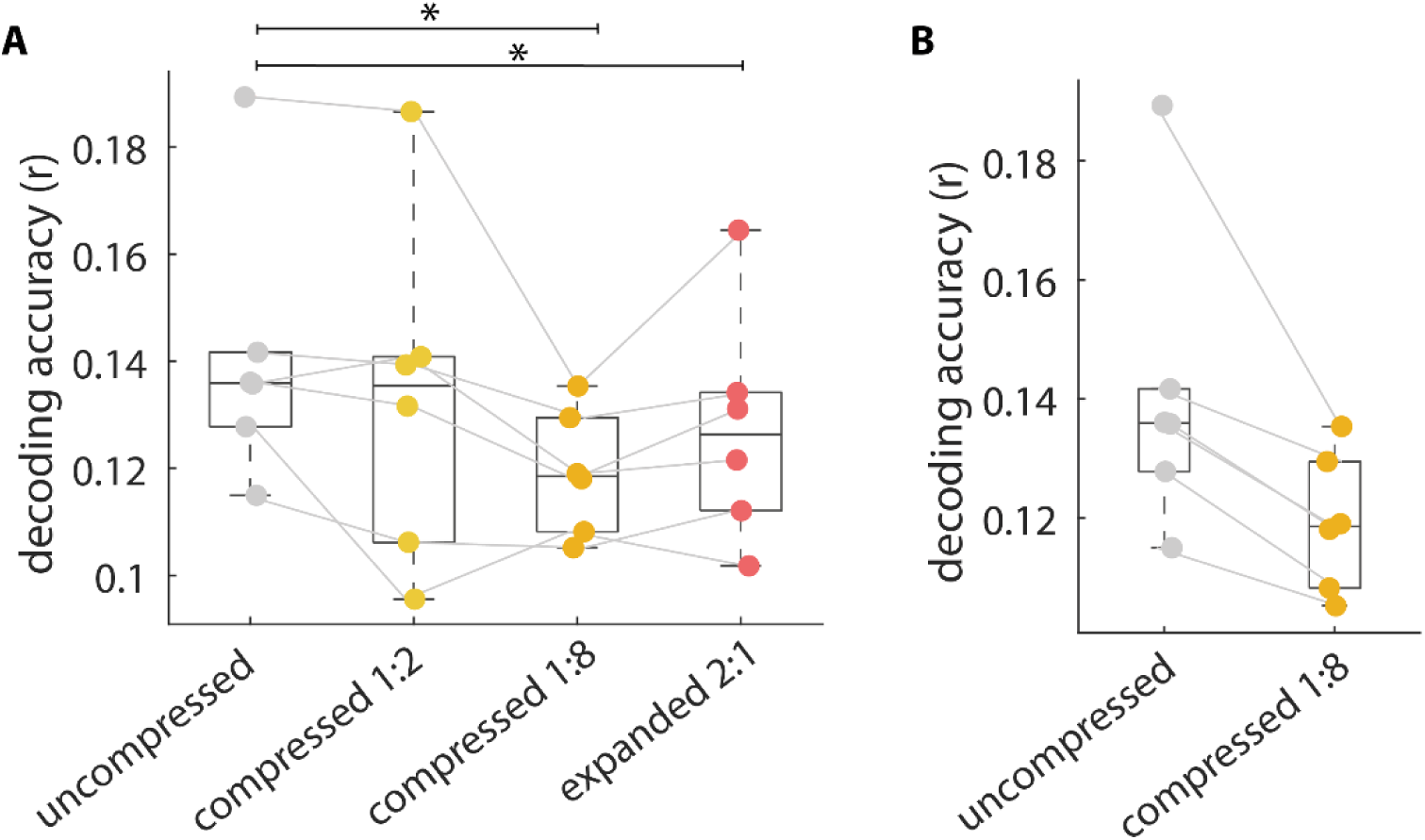
Pilot study: Neural results for different compression and expansion ratios. **A.** Decoding accuracy (0-500 ms) refers to the Pearson correlation between the stimulus onset envelope and the estimated onset envelope, using all EEG channels. The boxes represent the interquartile range (25th to 75th percentile) of the data, with the median indicated by a line inside the box, with dots representing individual subject data and connection lines indicating the same subjects. The whiskers extend to the most extreme data point, excluding outliers. **B** This plot shows the decoding accuracy for the uncompressed baseline compared to the 1:8 compression ratio, with dots representing individual subject data and connection lines indicating the same subjects. The boxes represent the interquartile range (25th to 75th percentile) of the data, with the median indicated by a line inside the box. The whiskers extend to the most extreme data point, excluding outliers.

In all our analysis, we used the onset envelopes of the actually presented signals. As a control, we also conducted an analysis using the uncompressed onset envelope, even when the signal was expanded or compressed, while keeping the ridge regression parameter λ constant. In this alternative analysis, we observed that most (5 out of 6) participants demonstrated a decrease in neural tracking, as measured by decoding accuracy, for compressed (1:8) speech.

Based on the results of the pilot experiment, we concluded that using a 1:8 compression ratio for amplitude compression reduces the brain’s ability to track speech. Therefore, we chose to use this compression ratio in our main experiment, which also included attention as an experimental factor. We made this decision for two main reasons. Firstly, our initial hypothesis that compression on ignored streams would increase neural separation only works when participants focus their attention on one stream and ignore the other. Secondly, attention could also have affected the compression effect in the pilot experiment, as participants may not have been attending to the compressed (1:8) speech stream.

### Amplitude compression impairs detection performance

We first analysed the behavioural data in terms of the proportion of detected repeats and response speed (Figure 4). For the main effects, we expected that attention (to-be-attend to the cued stream) has a positive effect on the dependent variables, which means that attention leads to an increased proportion of detected repeats and an increased response speed (inverse of response time). In contrast, compression should have a negative effect on behaviour, that is, a decreased proportion of detected repeats and a decreased response speed. In addition, we the effect of attention should depend on compression, with compression on the ignored stream increasing behavioural performance in contrast to the attentional pair where no compression is applied to the ignored stream.

**Figure 4.**
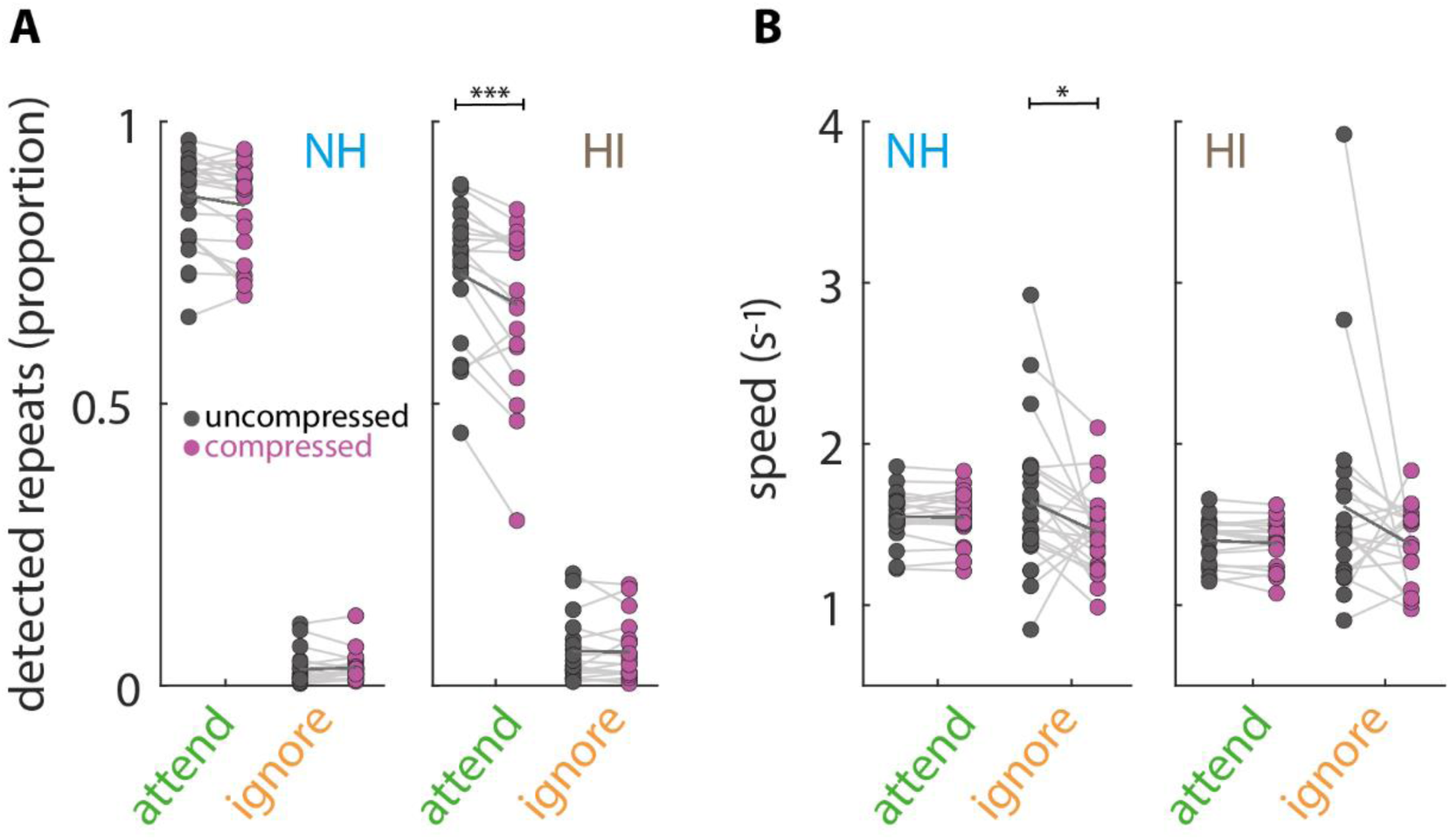
Proportion of detected repeats and corresponding response speed. **A** Left: Proportion of detected repeats per condition. Dots show individual NH (N=24, blue) and HI (N=19, brown) mean proportions of detected repeats. Light grey lines indicate the same subject. The bold line represents the mean of the group. Black dots indicate uncompressed tracking, while purple dots indicate compressed tracking. **B** Response speed per condition. Dots show individual (N=24, blue) and HI (N=19, brown) mean response speed. Light grey lines indicate the same subject. The bold line represents the mean of the group. Black dots indicate uncompressed tracking, while purple dots indicate compressed tracking.

Normal-hearing (NH) participants were well able to detect repeats in the attended uncompressed stream (Figure 4 A left; mean accuracy: 0.87, 95% CI: [0.84, 0.90]; mean speed: 1.56 s^−1^, 95% CI: [1.49, 1.62 s^−1^]). They were also well able to detect repeats in attended compressed stream (mean accuracy: 0.86, 95% CI: [0.82, 0.89]; mean speed: 1.55 s^−1^, 95% CI: [1.49, 1.62 s^−1^]). They made a only few false alarms for the ignored uncompressed stream (false alarm rate: 0.03, 95% CI: [0.02, 0.04]; mean speed: 1.65 s^−1^, 95% CI: [1.46, 1.84 s^−1^]) and for the ignored compressed stream (false alarm rate: 0.03, 95% CI: [0.02, 0.04]; mean speed: 1.44 s^−1^, 95% CI: [1.34, 1.55 s^−1^]). Jointly, the number of hits and false alarms indicate that participants were attending to the cued speech stream. We found no significant difference in mean accuracy (b = 0.02, SE = 0.06, OR = 1.02, 95% CI [0.99, 1.14], p = .74) between the compressed and uncompressed streams.

NH participants responded with similar response speed to repeats in attend and ignored speech, no significant differences were observed (Figure 4 B left; b = 0.036, SE = 0.05, t(12268) = 0.74, p = 0.46). We found a significant main effect of compression (b = 0.36, SE = 0.05, t(12260) = 7.36, p < .001), indicating that compression on speech streams led to a decreased response speed. However, this main effect was qualified by a significant interaction with attention (b = 0.69, SE = 0.1, t(12260) = 7.11, p < .001). A closer examination of the interaction via post hoc tests showed that the decrease in response speed on compressed speech was driven by ignoring (b = -0.7, SE = 0.1, t(12260) = -7.35, p < .001) not attending (b = -0.012, SE = 0.017, t(12260) = -0.7, p = 1). That is NH participants showed slower responses to false alarms in the ignore compressed stream.

Comparing the hit rate between the within attentional pairs of interest (attend uncompressed and ignore compressed vs. attend uncompressed and ignore uncompressed), we found no significant difference (b = -0.09, SE = 0.07, OR = 0.92, 95% CI [0.8, 1.01], p = .22) We found a similar pattern comparing the response speed. No significant differences between attend uncompressed and ignore compressed vs. attend uncompressed and ignore uncompressed (b = -0.012, SE = 0.01, t(12252) = -1.2, p = .24).

Participants with hearing impairment (HI) were as well able to detect repeated sounds in both the uncompressed and compressed attended streams, with mean accuracies of 0.73 (95% CI: [0.67, 0.79]) and 0.68 (95% CI: [0.61, 0.75]), respectively (Figure 4 A right). The difference in accuracy between the uncompressed and compressed attended streams was significant (Figure 4 A right, SE = 0.03, OR = 0.75, p < .001). HI participants also made very few false alarms in detecting repeated sounds in the ignored uncompressed and compressed streams, with mean accuracies of 0.06 (95% CI: [0.03, 0.09]) and 0.06 (95% CI: [0.03, 0.09]), respectively. The difference in false alarms between the compressed and uncompressed ignored streams was not significant (SE = 0.08, OR = 0.96, p = 1). We found no significant differences between attention (b = -0.02, SE = 0.02, t(8197) = 1.35, p =.176) and compression in the hearing impaired group (b = -0.02, SE = 0.02, t(8197) = 0.97, p =.334).

These results suggest that there is a significant difference in performance within the HI group between the uncompressed and compressed attended streams, with more accurate responses for the uncompressed attended stream.

### Compression decreases neural speech tracking in NH and HI participants

The strength of a speech stream’s representation in the EEG is reflected by neural tracking (see methods for details). Analysis of the neural tracking (0-500 ms) for normal hearing participants revealed significant main effects of attention , which means that attended speech is stronger tracked than ignored speech (b = 0.08, SE = 0.02, t(9185) = 3.9, p < .001) and compression, which means that uncompressed speech is stronger tracked than compressed speech (b = 0.13, SE = 0.02, t(9185) = 6.4, p < .001; Figure 5 A). Attention and compression had no significant interaction (b = -0.06, SE = 0.04, t(9185) = -1.36, p =.175).

**Figure 5.**
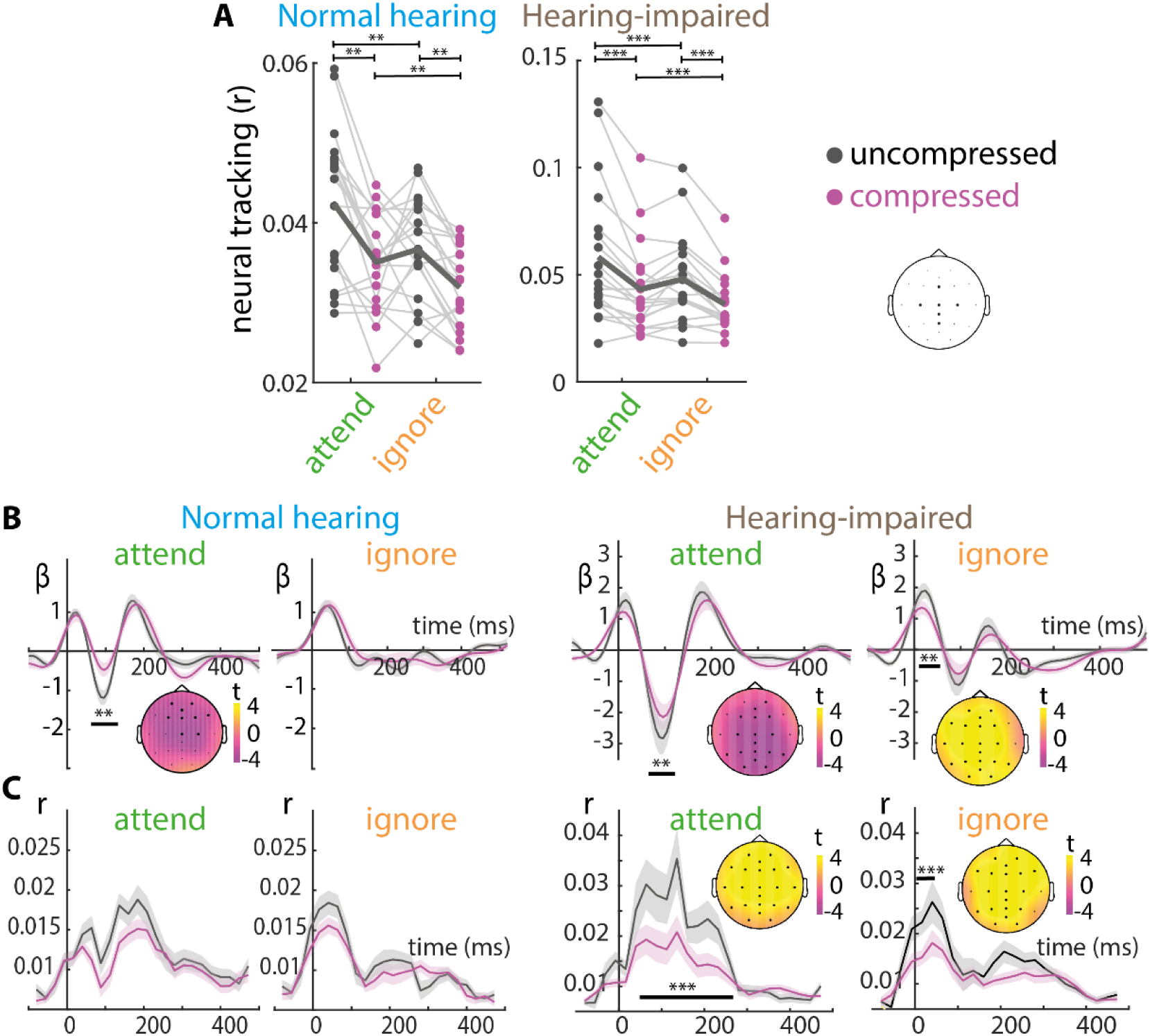
Neural tracking and temporal response functions (TRF). **A** Neural tracking (r) refers to the encoding accuracy (0-500 ms) obtained by multiplying estimated TRFs with the presented audio envelope. Line plot shows single-subject data (NH =24, HI = 19) averaged across channels of interest (C3, C4, Fz, Cz, Pz, CPz). Connection lines between dots indicate the same subject. Black dots indicate uncompressed tracking, while purple dots indicate compressed tracking. It’s important to note that the axes are scaled differently between the normal hearing and hearing-impaired plots. **B** TRF β-weights are averaged across subjects (NH = left, HI = right) and channels of interest. Inset displays the topographical distribution of t-values for the significant differences observed between conditions. Shaded areas depict the standard error for each time lag across subjects. **C** Unfolding neural tracking across time lags (-100-500 ms). Solid lines show the averaged neural tracking (r) across subjects and channels of interest (topographic map). Shaded areas show the standard error for each time lag across subjects. Inset displays the topographical distribution of t-values for the significant differences observed between conditions.

Analysis of the neural tracking (0-500 ms) for hearing-impaired participants revealed significant main effects of attention, which means that attended speech is stronger tracked than ignored speech (b = 0.008, SE = 0.001, t(7270) = 8.6, p < .001) and compression, which means that uncompressed speech is stronger tracked than compressed speech (b = 0.013, SE = 0.001, t(7270) = 13.5, p < .001; Figure 5 A). Attention and compression had no significant interaction (b = -0.003, SE = 0.001, t(7270) = -1.67, p =.094).

To assess the statistical significance of differences in TRFs between conditions for normal hearing participants, we used a cluster permutation test (Figure 5 B left). We found a significant negative difference comparing attended uncompressed and attended compressed TRFs (56-112 ms; cluster p-value = .003) which indicates a larger N1 amplitude for the attended uncompressed signal. No significant differencec between ignore uncompressed and ignore compressed were observed.

For hearing impaired participants (Figure 5 B right), we found a significant negative difference comparing attended uncompressed and attended compressed TRFs (64-112 ms; cluster p-value = .003) which indicates a larger N1 amplitude for the attended uncompressed signal. In addition, we found a significant difference comparing ignored uncompressed and ignored compressed TRFs (8-48 ms; cluster p-value = .006).

A cluster permutation test (Figure 5 C left) yielded no significant differences between attend uncompressed and attend compressed, or between ignore-uncompressed and ignore-compressed trials, respectively.

For hearing impaired participants (Figure 5 C right), we found a significant negative difference comparing attended uncompressed and attended compressed time-shifted neural tracking (40-256 ms; cluster p-value < .001) which indicates a smaller time-shifted neural tracking for the attended compressed signal. In addition, we found a very early significant difference of ignored uncompressed to ignored compressed TRFs (0-64 ms; cluster p-value < 001).

### Increased neural separation when ignored stream is compressed

Based on our second hypothesis that compression on the ignored stream increases the neural separation between the attended and ignored streams, we took a closer look on the attentional differences between attended uncompressed & ignore uncompressed vs. attended uncompressed & ignore compressed.

Analysis of the neural tracking (0-500 ms) for normal hearing participants within attentional pairs revealed (Figure 6 A, upper panel) a significant effect of attention when both streams were uncompressed (t(23)=2.35, Cohen’s d = 0.53, p =0.03). When only the ignored stream is compressed, we see significantly more neural tracking of the attended stream (t(23)=4.1, Cohen’s d = 0.9, p < .001).

**Figure 6.**
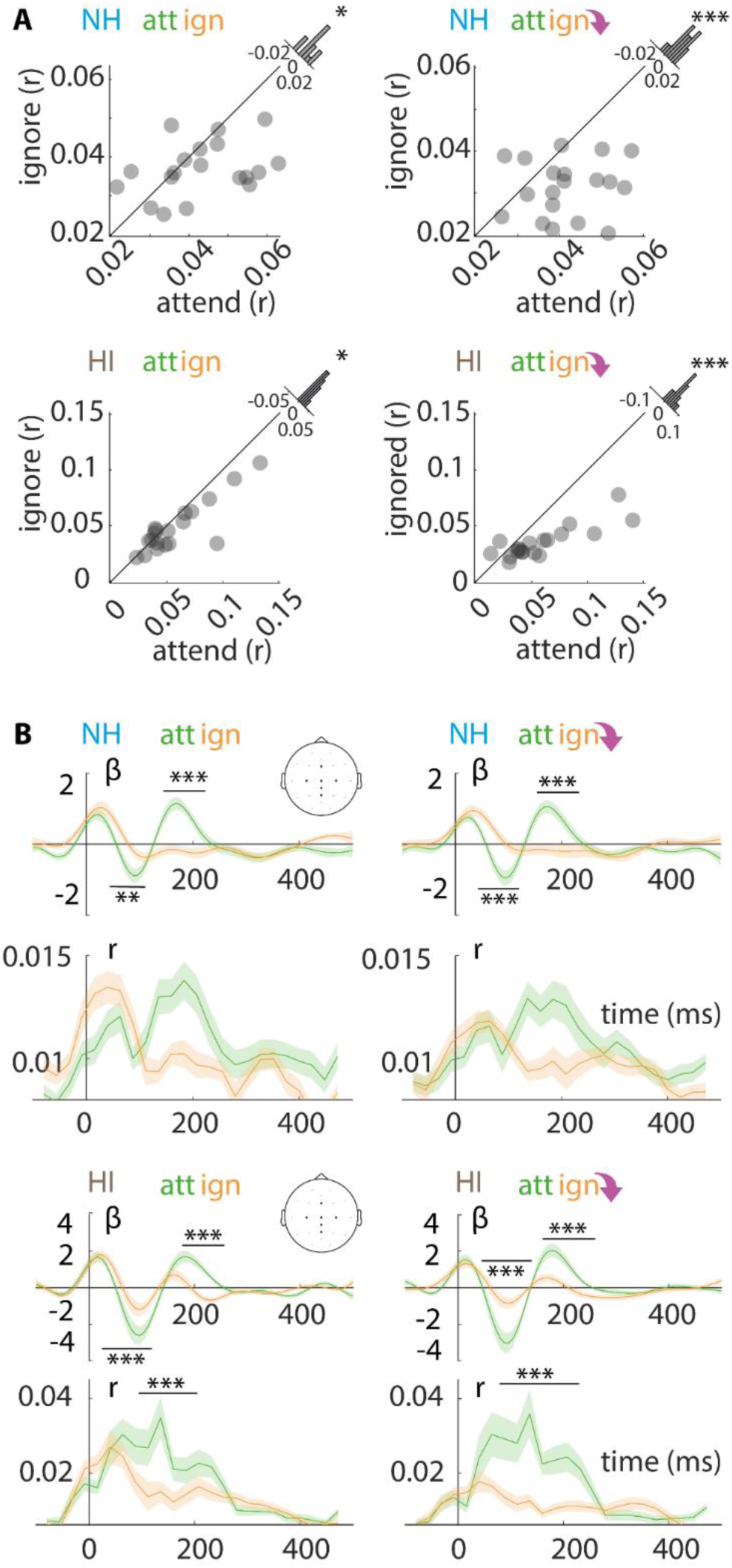
Neural tracking and TRF within attentional pairs for NH and HI participants. **A** Neural tracking (r) refers to the encoding accuracy (0-500 ms) based on estimated TRFs and envelopes. 45◦ plot shows single-subject data averaged across channels of interest for the attentional pairs. **B** Top: TRF β-weights and time-shifted encoding accuracy are averaged across subjects and channels of interest for the four attentional pairs. Shaded areas depict the standard error for each time lag across subjects. Black bars indicate significant differences between attend and ignore TRF. Bottom: Unfolding neural tracking across time lags (-100-500 ms). Solid lines show the averaged neural tracking (r) across subjects and channels of interest (topographic map). Shaded areas show the standard error for each time lag across subjects.

We conducted an analysis of neural tracking (0-500 ms) for hearing-impaired participants within attentional pairs (attend vs. ignore; Figure 6 A, lower panel) and found a significant difference between attend and ignore when both streams were uncompressed (t(18)=4.1, Cohen’s d = 0.7, p = 0.01). We did also observe a significant difference in the attentional pair when only the ignored stream was compressed (t(18)=4.3, Cohen’s d = 1, p < .001).

To assess the statistical significance of differences in TRFs within pairs for normal hearing participants, we used a cluster permutation test (Figure 6 B). We found one negative and one positive cluster (PC, NC) for both attentional pairs: attend uncompressed & ignore uncompressed (NC: 40-104 ms, cluster p-value = .002; PC: 136-232 ms, cluster p-value < .001), attend uncompressed & ignore compressed (NC: 40-112 ms, cluster p-value = < .001; PC: 144-248 ms, cluster p-value < .001). We found no significant differences in time-shifted neural tracking for normal hearing participants.

To assess the statistical significance of differences in TRFs within pairs for hearing-impaired participants, we used a cluster permutation test (Figure 6 B). We found one negative and one positive cluster (PC, NC) for both attentional pairs: attend uncompressed & ignore uncompressed (NC: 32-128 ms, cluster p-value < .001; PC: 160-248 ms, cluster p-value < .001), attend uncompressed & ignore compressed (NC: 48-136 ms, cluster p-value = < .001; PC: 144-304 ms, cluster p-value < .001).

For hearing impaired participants (Figure 5 C right), we found a significant difference for the attentional pair attend uncompressed & ignore uncompressed in time-shifted neural tracking (88-208 ms; cluster p-value < .001). We also found a significant difference for the attentional pair attend uncompressed & ignore compressed (64 -232 ms; cluster p-value < 001).

### Enhanced neural representation of uncompressed attended speech when the ignored stream is compressed in hearing-impaired old participants

We investigated the differences within the attentional pairs in more detail to determine whether the observed variations are due to differences in the uncompressed attended streams or within the compressed ignored streams. If the differences stem from the latter, it would suggest a potential attentional benefit of compressing the ignored streams.

Based on the PTA 25 dB HL (Fig. 7A) threshold we divided the cohort of age-matched older participants into two groups: those with normal hearing (PTA ≤ 25 dB HL, N =12) versus those with hearing impairment (PTA > 25 dB HL, N =19).

**Figure 7.**
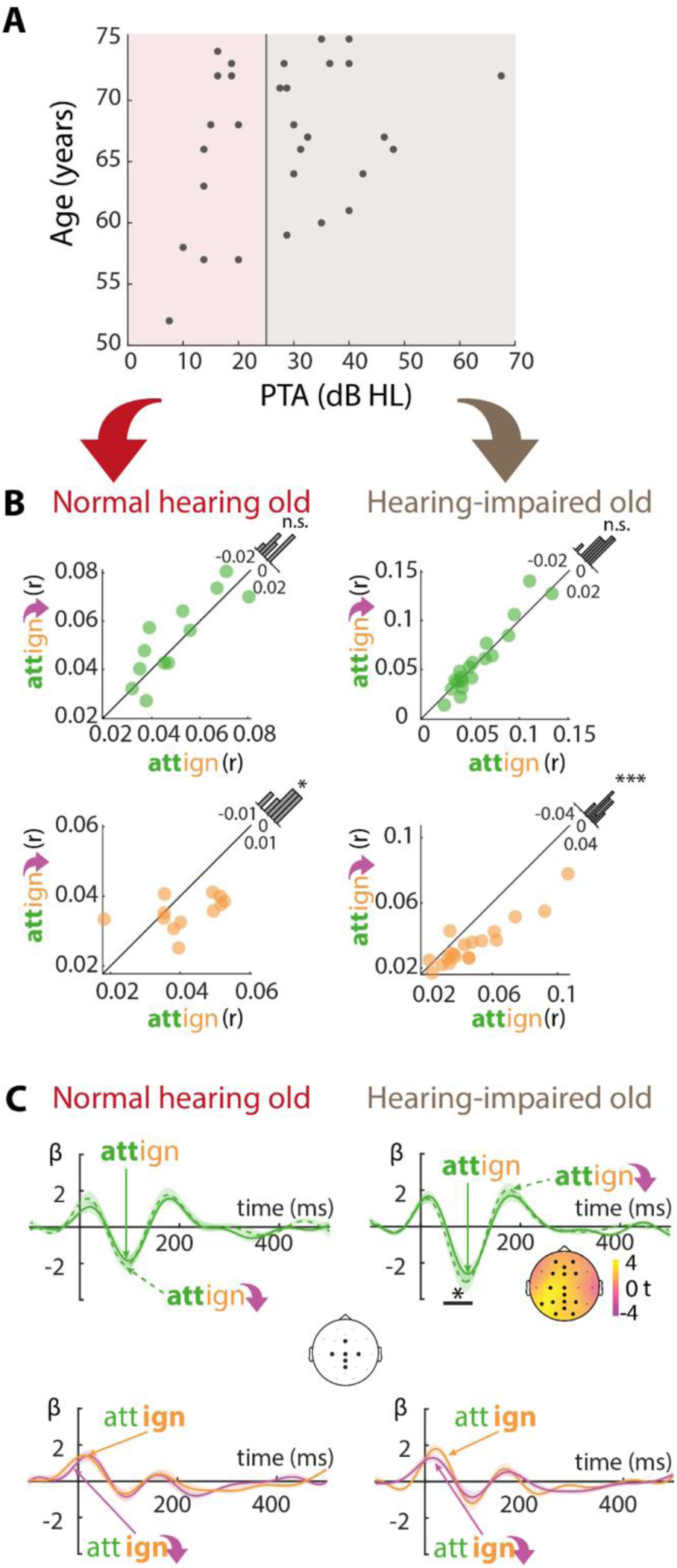
Comparison between attended and ignored neural tracking arising from different simultaneously presented attentional pairs for age-matched NH versus HI participants. **A** Relationship of Pure Tone Average (PTA) in decibels hearing level (dB HL; x axis) to age in years (y-axis; group of young, normal hearing not shown). Each point represents an individual’s PTA and age. Vertical line marks the threshold conventionally used to mark hearing loss: those with normal hearing (PTA ≤ 25 dB HL, N= 12) vs. those with hearing impairment (PTA > 25 dB HL, N=19). **B** Comparison between attended (upper panel) and ignored streams (lower panel) from attend and ignore uncompressed pairs vs. attend uncompressed and ignore compressed pairs for normal hearing old (left) and hearing-impaired old (right). Neural tracking (r) refers to the encoding accuracy (0-500 ms) based on estimated TRFs and envelopes. **C** TRF β-weights plots show the comparison between attended (upper panel) and ignored streams (lower panel) from attend and ignore uncompressed pairs vs. attend uncompressed and ignore compressed pairs for normal hearing old (left) and hearing-impaired old (right). Inset displays the topographical distribution of t-values for the significant differences observed between conditions.

In NH old participants, compressing (versus not compressing) the ignored stream yielded no statistically significant difference in the neural tracking of the attended uncompressed stream (t(11)=-1.087, Cohen’s d = -0.31, p =0.3, Fig. 7B). However, we found less neural tracking for the compressed ignored stream itself (t(11)=2.271, Cohen’s d = 0.66, p = 0.04) . We found also no significant differences comparing TRFs within the attentional pairs in the NH old group (Fig. 7C).

Hearing-impaired (HI) old participants showed also no difference in the neural tracking of the attended uncompressed stream when compressing (versus not compressing) the ignored stream (t(18)=-0.081, Cohen’s d = -0.02, p =0.9, Fig. 7B). Importantly, however, the TRF-derived N1 to the uncompressed attended speech stream in these HI listeners was larger when the ignored stream was compressed (compared to uncompressed; 48-104 ms, cluster p-value = 0.01). Note that this effect – compression in the ignored stream affecting the neural response to uncompressed attended speech – was specific to hearing-impaired listeners.

Fitting this effect of compression in the ignored stream, the HI old group also showed less neural tracking for the compressed (versus not compressed) ignored stream (t(18)=5.271 , Cohen’s d = 1.2, p < 0.001); hereby again matching the NH old group.

In NH young participants, there were no statistically significant differences in neural speech tracking between the attended uncompressed streams from both (t(23)=0.731, Cohen’s d = 0.17, p =0.474). We found a significant difference between the ignored uncompressed and ignored compressed stream (t(23)=3.954 , Cohen’s d = 0.9, p = 0.006) which indicates less neural tracking for the compressed ignored stream. We found also no significant differences comparing TRFs within the attentional pairs in the NH young group.

### Control analysis: Front–back location assignment does not confound neural and behavioural results

We considered the possibility that the front–back location assignment could have an indirect effect on our behavioural and neural measures. Between trials (and for some sustained trials), participants had to switch their attention between the front and back loudspeakers. We had randomized and balanced conditions across the two locations of the streams. Nevertheless, location as a factor was included in the statistical analysis to control for potential confounds.

We analysed the effects of location (front–back), attention, and compression on both behavioural performance and response time each in one model including all groups. Our results showed a significant main effect of location on behavioural performance (b = 0.25, SE = 0.03, OR = 1.2, 95% CI [1.2, 1.38], p < .001), indicating that participants detected more repeats in the front loudspeaker.

Importantly, however, there were no significant interactions between location and attention (b = -0.01, SE = 0.07,OR = 0.9, 95% CI [0.78, 1], p = 0.07), between location and compression (b = -0.08, SE = 0.07, OR = 0.92, 95% CI [0.8, 1.1], p = 0.2), or between location, compression, attention and age (b = -0.05, SE = 0.26, OR = 0.9, 95% CI [0.56, 1.57], p = 0.8).

Regarding response speed, we observed no significant main effect of location (b = 0.02, SE = 0.02, t = 1.34, p = 0.17) and no significant interactions between location and attention (b = -0.006, SE = 0.02, t = -0.25, p = 0.8), between location and compression (b = -0.01, SE = 0.02, t = -0.7, p = 0.47), and between location, compression, attention and age (b = -0.03, SE = 0.01, t = -0.3, p = 0.76).

Our neural analysis showed no significant main effect of location (b = 0.009, SE = 0.007, t = 1.16, p =0.25), and we found no significant interactions between location and attention (b = -0.001, SE = 0.001, t = -0.8, p = 0.43), between location and compression (b = 0.001, SE = 0.001, t = 0.09, p = 0.9), or between location, compression, attention and age (b = 0.0001, SE = 0.003, t = -0.001, p = 0.9). We thus consider it safe to conclude that the front-back location assignment did not confound the neural and behavioural measures.

### Control analysis: Auditory peripheral modelling of compressed and uncompressed speech

To avoid a potential confounding effect of how compressed speech gets processed along the ascending auditory pathway on our cortical neural results, we modelled the peripheral response to our stimulus material, employing a computational model of the human auditory periphery developed by Verhulst et al. (2018).

We used a mixed model to analyse the auditory nerve (AN) output (Figure 8A), with hearing status (NH vs. HI), signal manipulation (uncompressed vs. compressed), and frequency (500, 1000, 2000, and 4000 Hz) as fixed effects, and speech snippet as a random factor. Our analysis revealed a significant main effect of hearing status (b = 0.053, SE = 0.02, t = 2.4, p = 0.016), indicating higher log-transformed spike rates for audiograms matching our NH group compared to audiograms matching our HI group. We also observed a significant main effect of frequency (ref:1000 Hz; b = -1.3 to 0.9, SE = 0.03, t = -42 to 29, p < .001 for all), indicating higher log-transformed spike rates for higher frequencies. Importantly, there was no significant main effect on signal manipulation (b = 0.004, SE = 0.02, t = .2, p = 0.9) nor any significant interaction between frequency, signal manipulation and hearing overall(p=0.351). No other interactions were significant (all p > 0.05). Upon visual inspection the EFR (Figure 8 B), if at all, the compressed speech snippets appeared to lead to a higher amplitude in both simulations for normal hearing and hearing-impaired.

**Figure 8.**
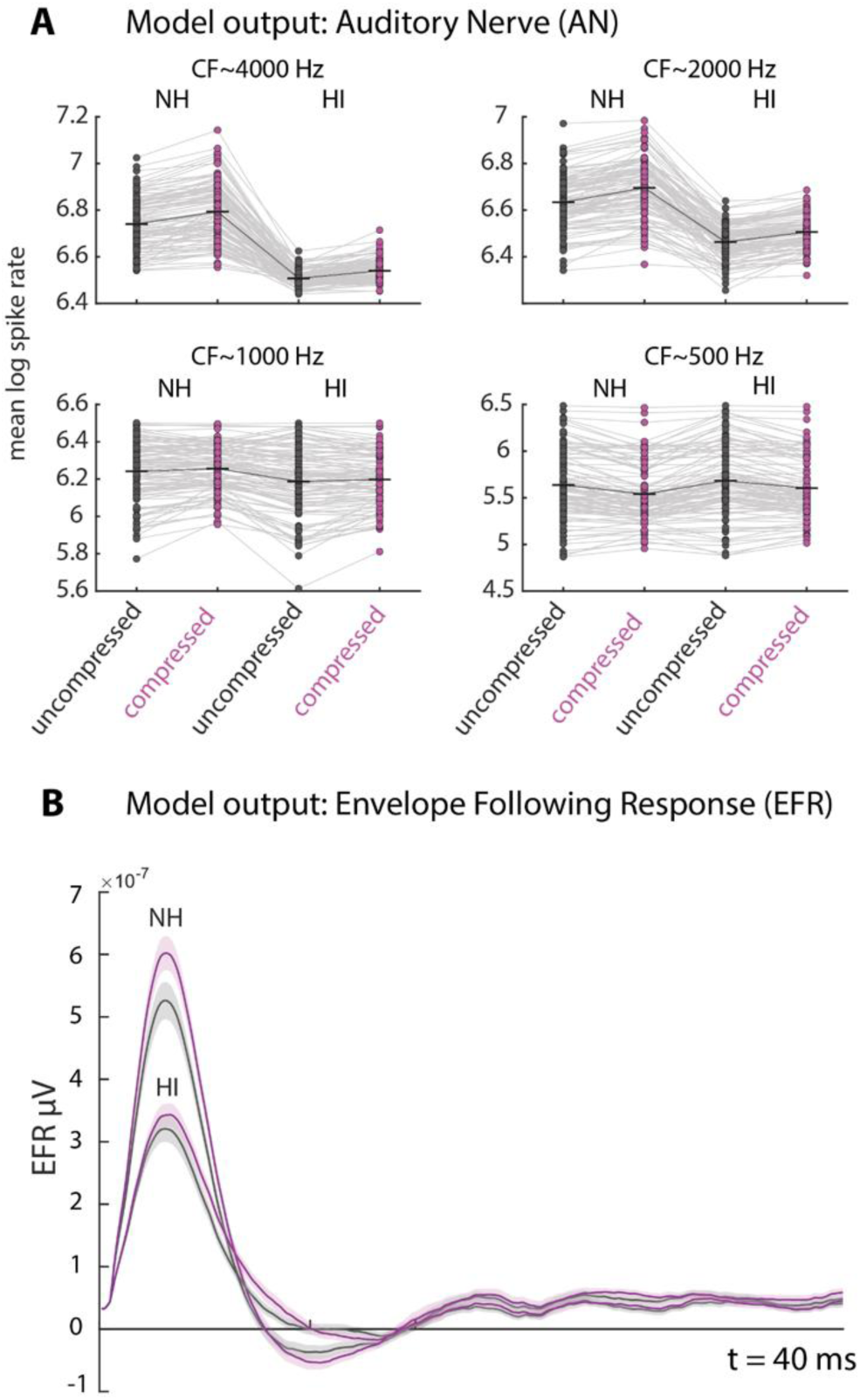
Simulated model output: AN and EFR. Panel **A** displays AN output for four center frequencies (500, 1000, 2000, and 4000 Hz), separated by normal hearing (NH) and hearing-impaired (HI) conditions and uncompressed (raw) and compressed (comp) speech snippets. Dots indicate different speech snippets. Connection lines indicate same snippet. Panel B shows the simulated EFR for NH and HI conditions, separate for compressed and uncompressed speech snippets. Shaded area shows SEM of different snippets, while solid line shows the mean across snippets. The colour purple indicates compression.

In sum, the peripheral modelling results indicate that effects of loudness-matched dynamic range compression along the auditory pathway are not able to account for the described neural tracking and cortical response effects described here.

## Discussion

In the present study, we aimed to investigate the effects of amplitude compression and its interactions with a listener’s selective attention goals on neural separation and listening behaviour. Studying normal and hearing-impaired participants using a psychophysically augmented continuous speech paradigm, our hypothesis was that compression decreases neural tracking. Furthermore, we expected that compression on ignored talkers could improve neural and behavioural markers of attentional separation.

### Amplitude compression on speech impairs behavioural performance

Hearing-impaired participants showed decreased behavioural responses when the attended streams were compressed. The findings are consistent with prior research indicating that fast-acting compression leads to reduced speech intelligibility (Drullman et al., 1994; Stone & Moore, 2004).

Two decades ago, Stone & Moore (2004, 2007) explored how compression reduces performance by introducing “across-source modulation correlation,” where compression causes signals to acquire a common modulation component, increasing perceptual fusion. This comodulation can make separating multiple speech signals challenging, as noted by Bregman (1994) and others. Our study mitigates this by applying amplitude modulation separately to each stream, ensuring that our results are not influenced by comodulation of the attended and ignored signals. Unlike comodulation, modulation reduction through compression is crucial in the present design. Our loudness matching pipeline’s dynamic range compressor adjusted gain based on intense signal components, typically corresponding to envelope peaks. Fast amplitude compression, employed in our pipeline, focuses on these peaks, reducing their gain. The pipeline enhances low-intensity signals, resulting in smaller amplitude modulation depth in compressed speech, distorting envelope fidelity (Stone & Moore, 1992). The non-instantaneous compressor operation can cause overshoots and undershoots, contributing to envelope distortion (Shetty, 2016; Verschuure et al., 1996).

When delving deeper into the repercussions of fast amplitude compression on speech signals, an intriguing facet emerges: the impact on the amplitude modulation depth and intensity contrast. Fast amplitude compression tends to reduce amplitude modulation depth and intensity contrast (Moore et al., 2003; Plomp, 1988). This is crucial for speech recognition as it primarily relies on temporal cues of speech ((Peelle et al., 2013; Shannon et al., 1995; Shetty, 2016; Stone & Moore, 2004). Furthermore, both speech streams experience reduced amplitude modulation, even though they are not comodulated. This results in a similar low amplitude modulation ratio for both streams, which can make it more challenging to separate them (Grimault et al., 2002).

### Amplitude compression decreases neural tracking of speech

To our knowledge, we have here presented the first study to explore the impact of amplitude compression on neural speech tracking using EEG recordings in both normal hearing and hearing-impaired individuals. Our findings revealed a significant effect of amplitude compression on neural tracking in normal hearing- and hearing-impaired participants.

Neural speech tracking relies on the temporal envelope of speech, as has been previously shown in various studies (Ding & Simon, 2012; Etard & Reichenbach, 2019; Kadir et al., 2019; Kerlin et al., 2010; Mesgarani & Chang, 2012; Obleser & Kayser, 2019; Rosen, 1992; Zion Golumbic et al., 2013). Therefore, it was expected that dynamic range compression, a form of signal processing that directly affects the temporal envelope of speech, might impair neural speech tracking. As described in more detail above, the fidelity of the temporal envelope is impaired (vowel to consonant ratio) due to reduction in amplitude modulation depth and overshoot and undershoots in amplitude compressed speech (Stone & Moore, 2007, 2007; Verschuure et al., 1996).

In the context of selective attention, our results showed a main effect of attention on neural tracking, with larger tracking of the attended speech compared to the ignored speech. This is in line with several previous studies that have shown enhanced neural responses to attended speech compared to ignored speech (e.g., Mesgarani & Chang, 2012; Zion Golumbic et al., 2013). We did not observe a significant interaction between compression and attention on neural tracking, suggesting that compression did not modulate the effect of selective attention on neural tracking in the first place.

Given the quasi-factorial nature of our experimental design, where attended and ignored speech streams are presented simultaneously, we also focused on attentional pairs, specifically examining the differences between the attended uncompressed and ignored uncompressed streams versus the attended uncompressed and ignored compressed streams. Our hypothesis was that compressing only the ignored stream would enhance neural separation between attended and ignored streams. We anticipated decreased neural tracking for the compressed, ignored stream and increased tracking for the attended stream, even compared to the attended stream when both streams were uncompressed. We found a significant difference in neural tracking between compressed and uncompressed ignored streams in both groups but no significant difference in tracking for uncompressed attended streams among normal-hearing participants. Most importantly, we observed a significantly larger N1 amplitude in the attended uncompressed TRF when the ignored stream was compressed, compared to when the ignored stream was uncompressed, but this effect was observed only in hearing-impaired participants and not in age-matched normal hearing controls.

### N1 amplitude increase in hearing-impaired participants’ uncompressed attended TRF when the ignored stream is compressed

Comparing hearing-impaired and normal hearing participants, we observed larger differential tracking and neural responses in hearing-impaired individuals between the attended and ignored streams. Normal hearing participants performed well (mean accuracy: 0.87), and the compression manipulation on the ignored stream might not have triggered the need for distractor suppression or target enhancement. In contrast, hearing-impaired participants tended to perform worse overall, potentially benefiting more from a compressed ignored talker. Hearing-impaired (HI) are more affected by compression than those with normal hearing participants. This is evident in all our neural measures including temporal response functions and neural speech tracking, especially in the comparison between compressed and uncompressed ignored streams, which showed weaker tracking for the compressed stream, suggesting additional suppression. This could facilitate the processing of the attended stream, leading to increased N1 TRF component of the uncompressed attended TRF when the ignored stream is compressed. However, to cleanly distinguish between target enhancement and distractor suppression, an additional baseline would be beneficial (Orf et al., 2023; Wöstmann et al., 2022).

For normal hearing participants, these findings endorse the idea that compression diminishes neural speech processing, leading to decreased neural tracking. The primary factor driving this effect is likely the distortion of envelope fidelity, as previously discussed. However, for hearing-impaired participants, there may be more to uncover. Future studies should explore whether hearing-impaired individuals might also experience benefits from more pronounced compression applied to ignored streams.

### Limitations

Despite out best efforts, the subcortical and cortical effects of dynamic range compression, especially in combination with counteracting adjustments for perceived loudness, are not trivial to capture: Using a computational model for auditory peripheral modelling, specifically focusing on the auditory nerve and the envelope following response, has its limitations compared to real measurements. Computational models inherently simplify the intricate biological processes of the auditory system. While such models aim to replicate physiological responses, they may not fully capture the complexity and variability of real neural responses. Limitations include assumptions made in modelling, potential oversimplifications of neural processing, and uncertainties in translating model outputs to real-world neural responses. Moreover, individual differences in the auditory system among participants may not be fully accounted for in a computational model. For a more comprehensive understanding, incorporating real measurements of the auditory periphery in future studies would enhance the validity and applicability of findings to actual neural responses in individuals.

## Conclusions

We here explored whether amplitude compression aids or hinders attentional neural speech tracking, for normal hearing as well as hearing-impaired participants. For normal hearing participants, applying amplitude compression to both streams hampers neural speech tracking, and no benefit from compression applied solely to the ignored stream is evident. While these findings generally extend to hearing-impaired participants, they exhibit a curious benefit in attentional separation in terms of neural speech tracking when the ignored speech stream is amplitude compressed. Using modern hearing aid algorithms that allow varying compression ratios for different spatial sources or investigating multiple distractors that become comodulated and fuse as one distracting source, could provide valuable insights into the psychoacoustic and neurophysiological effects of amplitude compression and its potential benefits.

## Acknowledgements

Research was supported by a research grant from WS Audiology, Erlangen, Germany, to JO.

